# Mitochondrial dysfunction and impaired osteogenic capacity define stress-induced osteoblast senescence

**DOI:** 10.64898/2026.09.13.751251

**Authors:** Tanja Frey, Hannah Vogg, Mubashir Ahmad, Yuta Nakano, Mona Vogel, Sandra Nungeß, Astrid Schoppa, Hartmut Geiger, Antia Ignatius, Jana Riegger

**Author notes:** Author email addresses: Tanja Frey; Hannah Vogg; Mubashir Ahmad; Yuta Nakano; Mona Vogel; Sandra Nungeß:; Astrid Schoppa; Hartmut Geiger; Anita Ignatius; Jana Riegger. **Institutional Review Board Statement** The study was conducted according to the guidelines of the Declaration of Helsinki and approved by the Ethical Committee of the University of Ulm, Germany (ethical approval number 76/25 and 265/22). **Informed Consent Statement** Informed consent was obtained from all subjects involved in the study.

## Abstract

Cellular senescence has emerged as a key contributor to age-related skeletal deterioration; however, the defining characteristics of senescent osteoblasts remain incompletely understood, hindering efforts to identify the cellular mechanisms that drive age-associated bone loss and potential therapeutic targets.

Here, we compared doxorubicin- and hydrogen peroxide-induced senescence and established a robust *in vitro* osteoblast model that enables the stable maintenance of stress-induced premature senescence.

Doxorubicin-treated MC3T3-E1 cells exhibited persistent growth arrest, a pronounced senescence-associated secretory phenotype (SASP), and impaired osteogenic function accompanied by marked mitochondrial dysfunction, including reduced respiratory capacity and altered mitochondrial morphology. Transcriptomic comparison with aged murine bone revealed a partial overlap with *in vivo* aging-associated gene expression signatures, particularly among genes involved in extracellular matrix organization and skeletal development, supporting the physiological relevance of the model. Importantly, key features of senescence, including the senescence-associated mitochondrial phenotype and impaired osteogenic capacity, were recapitulated in primary human osteoblasts.

Collectively, these findings establish a robust model of stress-induced osteoblast senescence and demonstrate that senescent osteoblasts exhibit mitochondrial dysfunction, impaired osteogenic capacity, and molecular features that resemble aspects of skeletal aging.

## 1. Introduction

Osteoporosis is a systemic skeletal disease characterized by reduced bone mass and deterioration of bone microarchitecture, resulting in increased bone fragility and fracture risk. It affects approximately one in three women and one in five men over the age of 50 (Das, Kotwal, Thakur, T, & Kumar, 2025). The disease is primarily driven by an imbalance in bone remodeling in which bone-resorption by osteoclasts exceeds and bone-formation by osteoblasts, due to impaired osteoblast differentiation and increased osteoclast activity (Pignolo, Law, & Chandra, 2021). This imbalance progressively worsens with advancing age, leading to continuous bone loss and increased skeletal fragility.

Although endocrine alterations contribute to osteoporosis progression, accumulating evidence indicates that intrinsic aging mechanisms within the bone microenvironment are major driver of age-related bone loss (Farr & Khosla, 2015). Among these, cellular senescence has emerged as a fundamental mechanism linking aging to impaired tissue homeostasis and the development of age-related diseases, including osteoporosis (Riegger, Schoppa, Ruths, Haffner-Luntzer, & Ignatius, 2023).

Cellular senescence is defined as a state of stable and largely irreversible state of cell cycle arrest induced by various cellular stressors, including DNA damage, oxidative stress, mitochondrial dysfunction, and telomere shortening (Campisi & d’Adda di Fagagna, 2007). While senescence initially acts as a beneficial tumor-suppressive mechanism by preventing the proliferation of damages cells, senescent cells accumulate with age, owing the persistent cellular stress and declining immune-mediated clearance. Their accumulation contributes to chronic inflammation, impaired tissue regeneration, and progressive functional decline (Collado, Blasco, & Serrano, 2007; van Deursen, 2014).

Besides replicative senescence caused by telomere attrition, senescence can also be induced independently of replicative exhaustion through external stressors, a process referred to as stress-induced premature senescence (SIPS). SIPS occur in both proliferating and post-mitotic cells following exposure to oxidative stress, genotoxic agents, mitochondrial dysfunction (Rossi et al., 2025), or chemotherapy-induced DNA damage (Lozano-Torres et al., 2019), and has been implicated in tissue injury, impaired regeneration, and numerous age-related diseases.

Mitochondrial dysfunction has emerged as a central hallmark of both aging and cellular senescence. Senescent cells exhibit characteristic mitochondrial alterations, including disrupted mitochondrial morphology, reduced respiratory function loss of membrane potential, increased reactive oxygen species (ROS) production and metabolic reprogramming. Together, these features constitute the senescence-associated mitochondrial phenotype (SAMP) (Martini & Passos, 2023). In bone tissue, mitochondrial dysfunction has been associated with impaired osteoblast differentiation, reduced bone-forming capacity, and the acquisition of senescence-associated characteristics (Schoppa et al., 2022). Damaged mitochondria promote excessive ROS production, which can initiate and reinforce SIPS. Conversely, the SAMP sustains the senescent phenotype by maintaining persistent DNA damage responses, enhance SASP expression, enforcing stable cell cycle arrest, and promoting resistance to apoptosis (Hu et al., 2022).

Despite growing evidence implicating osteoblast senescence in age-related bone loss, robust and physiologically relevant *in vitro* models that faithfully recapitulate the molecular and functional characteristics of senescent osteoblasts remain limited. In particular, the contribution of the SAMP to osteoblast dysfunction has not been comprehensively defined. Here, we establish a robust model of SIPS in osteoblasts and provide an in-depth characterization of its molecular, metabolic, and functional features, with a particular emphasis on mitochondrial dysfunction. Furthermore, by comparing transcriptomic signatures with aged murine bone and validating key findings in primary human osteoblasts, we assess the physiological relevance of this model as a platform for investigating osteoblast senescence and its potential contributions to age-related skeletal decline.

## 2. Material and Methods

### 2.1 Cultivation of pre-osteoblastic MC3T3-E1 cell line

Pre-osteoblastic MC3T3-E1 cells were cultivated in α-minimum essential medium (α-MEM; Gibco^TM^ by life technologies, Paisley, UK) supplemented with 10% fetal bovine serum (FBS), 2 mmol/L L-glutamine, and 100 U /100 µg /mL penicillin / streptomycin (all from PAN-Biotech, Aidenbach, Germany). Cells were maintained at 37°C, 95% humidity, and 5% CO_2_. Culture medium was replaced twice weekly, and cells were maintained at 70-80% confluency.

### 2.2 Isolation and cultivation of human osteoblasts

Human osteoblasts (hOB) were isolated as previously described (Hengartner, Fiedler, Ignatius, & Brenner, 2013). Briefly, cancellous bone tissue was obtained from the posterior tibial plateau during total knee replacement surgery following informed patient consent and approval by the Ethics Committee of the University of Ulm (Ethics Votum: 76/25 and 265/22). Approximately 5 g of cancellous bone was harvested, washed twice with phosphate-buffered saline (PBS; Gibco^TM^), and incubated in 0.05% collagenase (Sigma-Adrich, St. Louis, MO, USA) prepared in Dulbecco’s Modified Eagle Medium (DMEM (1 g/L D-Glucose); Gibco^TM^) supplemented with L-glutamine and pyruvate for 2 h at 37°C under rotation. Following enzymatic digestion, bone fragments were washed with DMEM and cultured in 6-well plates containing Ham’s F-12 medium supplemented with 10% FBS, 2 mmol/L L-glutamine, 100 U /100 µg /mL penicillin / streptomycin, and 2.5 μg/L amphotericin B (all from PAN-Biotech). Cells were maintained at 37°C in 95% humidity and 5% CO_2_. Culture medium was changed three times weekly. Following outgrowth, cells were maintained in DMEM supplemented with 10% FBS, 2 mmol/l L-glutamine, and 100 U /100 µg /mL penicillin / streptomycin (all from PAN-Biotech) and passaged no more than eight times.

### 2.3 Induction of cellular senescence using doxorubicin and hydrogen peroxide

Senescence of hOB and MC3T3-E1 cells was induced by using doxorubicin based on the protocol previously published by Kirsch et al (Kirsch, Ramge, Schoppa, Ignatius, & Riegger, 2022). Briefly, hOB and MC3T3-E1 cells were seeded at densities of 6000 cells/cm^2^ and 5000 cells/cm^2^, respectively, in their corresponding basal media. To induce a stable senescent phenotype, cells were treated with 0.1 μM doxorubicin (Doxo; Selleckchem, Houston, TX, USA) for five consecutive days. On day 5, 0.2 µM doxorubicin was administered. For oxidative stress-induced senescence, cells were treated with 100 μM hydrogen peroxide (H_2_O_2_; Sigma-Aldrich) twice, beginning one day after seeding, with a 24 h recovery period between treatments. A schematic overview of the stimulation protocols is provided in the Supplementary Figure 1A.

### 2.4 Alamar Blue Assay

Cell proliferation and cytotoxicity were assessed using a resazurin-based Alamar Blue assay. A 100x stock solution was prepared by dissolving 0.05 g resazurin sodium salt (Sigma-Aldrich) in 10 mL PBS (Gibco^TM^), followed by sterile filtration. The working solution was prepared by diluting the stock solution in 1:50 in PBS. For measurements, 50 μL resazurin working solution was added to 450 μL culture medium per well. Cells were incubated for 2-3 h at 37°C, and fluorescence was measured at an excitation wavelength of 550 nm and an emission wavelength of 590 nm in an Infinite M200 Pro microplate reader (Tecan Deutschland GmbH, Crailsheim, Germany) with i-control software version 2.0 (Tecan Deutschland GmbH).

### 2.5 Immunofluorescence Staining for Ki-67 and γ-H2A.X

For immunofluorescence staining, MC3T3-E1 cells and hOB were seeded at a density of 23,500 cells/cm^2^.

#### Ki-67 staining

Cells were fixed with 4% paraformaldehyde (PFA; Invitrogen, OR, USA), permeabilized with 0.1% Triton-X-100 (Merck KGaA, Darmstadt, Germany), and blocked using protein blocking buffer (Abcam, Boston, MA, USA) for 1 h at 37°C. Cells were incubated overnight at 4°C with anti-Ki-67 primary antibody (rabbit monoclonal, ab16667, Abcam; 1:250 dilution) prepared in antibody diluent (AB64211, Abcam). Subsequently, cells were incubated with Alexa Fluor^TM^ 488 goat anti-rabbit secondary antibody (ab150077, Abcam; 1:200 dilution) for 30 min at room temperature protected from light. Nuclei were counterstained using NucBlue^TM^ (Invitrogen) according to the manufacturer’s instructions for 5 min. Slides were mounted using FluorSave^TM^ Reagent (EMD Millipore Corp., USA), and images were acquired using an Axiovert 35 microscope and AxioVision software version 4.8.2 (Zeiss, Oberkochen, Germany).

#### γ-H2A.X staining

For detection of DNA double-strand breaks, cells were fixed with 4% PFA and permeabilized with 0.5% Triton X-100 for 10 min at room temperature. After washing with 0.05% Tween-20 (Merck KGaA), cells were blocked for 1 h at room temperature. Cells were incubated overnight at 4°C with anti-phospho-H2A.X (Ser139) antibodies (05-636, mouse monoclonal JBW301, Millipore, 1:200 for hOB; 83307-2-RR, rabbit monoclonal 5N19, Proteintech, 1:200 for MC3T3-E1). Following washing steps, cells were incubated with species-appropriate secondary antibodies (ab175473, Alexa Fluor 568 goat anti-mouse for hOB; Alexa Fluor goat anti-rabbit for MC3T3-E1; 1:200) for 30 min 37°C. Nuclei were counter stained with NucBlue^TM^ (Invitrogen) for 5 min. Slides were mounted with FluorSave^TM^ and imaged using Axiovert 35 (Zeiss) and corresponding software as well as BZ-X1000 and BZ-X Series Application version 1.0.2.89 (KEYENCE DEUTSCHLAND GmbH, Frankfurt am Main, Germany) in DAPI, FITC or TexasRed.

### 2.6 JC-1 Assay

Mitochondrial membrane potential in MC3T3-E1 cells was analyzed by using JC-1 dye. Briefly, cells were detached using Trypsin/EDTA (PAN-Biotech), and at least 500,000 cells were used per sample. Cells were washed with PBS (Gibco^TM^) and incubated with 1 μM JC-1 dye (Hycultec GmbH, Beutelsbach, Germany) for 30 min at 37°C. For the positive control 50 μM FCCP (Selleckchem) was added after 25 min of incubation and incubated for the remaining 5 min. Subsequently, cells were washed twice with PBS and resuspended in PBS for flow cytometric analysis. At least 50,000 events per sample were recorded using FACSCalibur flow cytometer equipped with dual-laser technology and analyzed using CellQuest software version 5.2.1 (BD Biosciences, Franklin Lakes, NJ, USA). Mitochondrial membrane potential was determined by calculating the red-to-green fluorescence intensity ratio. A representative gating strategy is provided in the Supplementary Figure 1B.

### 2.7 Cell Cycle Analysis

Cell cycle distribution of hOB and MC3T3-E1 cells was analyzed using Vybrant^TM^ DyeCycle^TM^ Green stain (Invitrogen, Carlsbad, CA, USA) according to the manufacturer’s instructions. A minimum of 250,000 events per sample was analyzed using FACSCalibur flow cytometer (BD Biosciences) with CellQuest software version 5.2.1. Representative gating strategies are shown in the Supplementary Figure 1C.

### 2.8 MitoSOX Assay

Mitochondrial superoxide accumulation in MC3T3-E1 cells was assessed using MitoSOX^TM^ Red (Thermo Fisher Scientific, Waltham, MA, USA). Briefly, 250,000 cells were washed twice with pre-warmed PBS and incubated with 2 μM MitoSOX^TM^ Red working solution for 15 min at 37°C protected from light. Following incubation, cells were washed three times with warm PBS and resuspended in PBS for flow cytometric analysis. Unstained control cells were included to determine autofluorescence, which was subsequently subtracted from MitoSOX^TM^ fluorescence measurements. A minimum of 10,000 events per sample was analyzed using BD Fortessa SORP with FACS DIVA 8.0 (BD Biosciences).

### 2.9 Senescence-Associated β-Galactosidase (SA-β-gal) Staining

SA-β-gal staining was performed using a Senescence β-Galactosidase Staining Kit (Cell Signaling Technology, Danvers, MA, USA) as previously described (Kirsch et al., 2022). In case of crystal formation, slides were briefly rinsed with dimethyl sulfoxide (DMSO; AppliChem GmbH, Darmstadt, Germany).

### 2.10 Mitochondrial stress test

Doxorubicin-treated and untreated MC3T3-E1 cells were seeded in a Seahorse XFe96/XF Pro Cell culture Microplate at a density of 4000 cells per well for doxorubicin-treated MC3T3-E1 and 3000 cells per well for untreated control. Doxorubicin-treated and untreated hOB were seeded with a density of 7500 cell per well for both conditions. On the day of measurement, the cells were incubated for 1 h in bicarbonate-free DMEM containing 5 mM HEPES, 10 mM glucose, 1 mM pyruvate, 2 mM glutamine. Oxygen consumption and extracellular acidyfication rates (OCRs and ECARs) were measured simultaneously using a Seahorse XFe96 Flux Analyzer (Agilent Technologies). Uncoupled (proton leak) respiration was profiled by injecting 1.5 µM oligomycin (an ATP synthase inhibitor) and full substrate oxidation capacity was determined by injecting 2 µM (hOB) or 2.5 µM (MC3T3-E1) carbonylcyanide-p-trifluoromethoxyphenylhydrazone (FCCP, a chemical uncoupler). Non-mitochondrial respiration was determined by injecting 0.5 µM antimycin A and 0.5 µM rotenone (ETC inhibitors). Data were normalized to by JanusGreen staining (Raspotnig et al., 1999). Mitochondrial ATP and glycolytic ATP production rates were calculated according to Desousa et al. (Desousa et al., 2023).

### 2.11 Visualization of mitochondria using transmission electron microscopy

Doxorubicin-treated and untreated MC3T3-E1 and hOB were seeded with a density of 21,000 cells/cm^2^ for doxorubicin-treated cells respectively and were vitrified through high-pressure freezing on carbon coated sapphire discs in a Wohlwend HPF compact 01 high-pressure freezer. Freeze-substitution was performed with 0.2% (v/v) osmium tetroxide, 0.1% (w/v) uranyl acetate and 5% (v/v) H_2_O in acetone by gradually increasing the temperature over 17 h from -90°C to 0°C. Temperature was kept at 0°C for 1 h and then increased within 1 h to room temperature. Samples were washed with acetone, followed by embedding in Epon 812 in hour consecutive steps at room temperature: 33% Epon 812/67% acetone for 1h, followed by 67% Epon 812/33% acetone for 3h, 90% Epon 812/10% acetone overnight, and pure Epon 812 overnight (v/v). Samples were transferred into fresh Epon 812 and polymerized at 60°C for 72 h. Embedded cells were sectioned in ∼70 nm Steps with a Leica EM UC7 ultramicrotome. Sections were captured on a formvar film on glow-discharged copper grids. Images were acquired with a 120 kV Jeol JEM-1400 Transmission electron microscope and a Celeta CCD camera (Olympus).

### 2.12 Osteogenic differentiation

To assess osteogenic differentiation potential, untreated and doxorubicin-treated MC3T3-E1 cells and hOB were cultured under osteogenic conditions. MC3T3-E1 cells were seeded at a density of 52,600 cells/cm^2^, whereas hOB were seeded at a density of 21,000 cells/cm^2^. Osteogenic differentiation was induced using DMEM (Gibco^TM^) supplemented with 10% FBS, 2 mmol/L L-glutamine, 100 U /100 µg /mL penicillin / streptomycin (all from PAN-Biotech), 0.1 μM dexamethasone, 10 mM β-glycerophosphate disodium salt hydrate, and 2 mM L-ascorbic acid-2-phosphate (Sigma-Aldrich). Medium was replaced three times weekly. Osteogenic differentiation of MC3T3-E1 cells was evaluated on day 7, day 14 and day 21 using qualitative and quantitative alkaline phosphatase (ALPL) staining and Alizarin Red S staining respectively.

### 2.13 ALPL and Alizarin Red S Staining

Prior to ALPL and Alizarin Red S staining, cell viability was assessed using the Alamar Blue assay as described above to later normalize either ALPL activity or Alizarin Red S to it.

#### ALPL staining

For qualitative ALPL staining on day 7, the Leukocyte Alkaline Phosphatase Kit (Sigma-Aldrich) was used. Although the kit is marketed for leukocytes/lymphocyte-related application, it produced positive staining in fixed adherent cells under our conditions. Cells were washed twice with PBS (Gibco^TM^) and fixed with 4% PFA (Thermo Fisher Scientific) for 10 min at room temperature. The staining solution consisted of 93.76% distilled water, 2.08% sodium nitrate, 2.08% Fast Red Violet (FRV), and 2.08% naphthol. Sodium nitrate and FRV were mixed first and incubated for 2 min prior to the addition of distilled water and naphthol. Cells were incubated with staining solution for 1 h at room temperature protected from light. Subsequently, cells were washed with PBS, and whole-well scans were acquired using a BZ-X1000 microscope (Keyence Deutschland GmbH). ALPL activity was additionally quantified using the Amplite^TM^ Colorimetric Alkaline Phosphatase Assay Kit (AAT Bioquest®, Sunnyvale, CA, USA) according to manufacturer’s instructions on day 7. Absorbance was measured at 405 nm with an Infinite M200 Pro microplate reader (Tecan Deutschland GmbH, Crailsheim, Germany) with i-control software version 2.0 (Tecan Deutschland GmbH).

#### Alizarin Red S staining

To quantify matrix mineralization, Alizarin Red S staining was performed at day 14 and day 21. Cells were fixed with ice-cold 70% ethanol for 1 h at room temperature. Subsequently, cells were stained with 40mM pH 4.2 Alizarin Red S solution (Sigma-Aldrich) for 10 min under gentle agitation. After staining, cells were washed with distilled water, and whole-well scans were obtained using the BZ-X1000 imaging system (Keyence Deutschland GmbH). For quantitative analysis, 10% hexadecylpyridinium chloride (Sigma-Aldrich) was added and incubated for 15 min under agitation to dissolve bound stain. Absorbance was measured at 652 nm with an Infinite M200 Pro microplate reader (Tecan Deutschland GmbH, Crailsheim, Germany) with i-control software version 2.0 (Tecan Deutschland GmbH).

### 2.14 Quantitative Real-Time PCR (qRT-PCR)

Total RNA was isolated from at least 100,000 MC3T3-E1 cells or hOB using the ReliaPrep^TM^ RNA Miniprep System (Promega, Madison, WI, USA). Reverse transcription was performed using SMARTScribe^TM^ Reverse Transcriptase (Takara Bio USA, San Jose, CA, USA). Quantitative real-time PCR was performed using a StepOnePlus^TM^ Real-Time PCR system (Applied Biosystems, Darmstadt, Germany). Relative gene expression was calculated using the ΔΔCt method.

*GAPDH/Gapdh*, *HPRT1/Hprt1* and *TMEM199* served as housekeeping genes as previously described (Hernandez-Segura, Rubingh, & Demaria, 2019). TaqMan^TM^ Gene Expression Assays (Thermo Fisher Scientific) used in this study included:

#### Human

*CDKN1A* (Hs00355782_m1; *CDKN2A* (Hs00923894_m1); *GAPDH* (Hs02758991_g1); *HPRT1* (Hs02800695_m1); *IL1β* (Hs00174097_m1); *IL6* (Hs00174131_m1); *IL8* (Hs01555410_m1); *TMEM199* (Hs01022209_m1); *TNF* (Hs01113624_g1); *TNFRSF11B* (*OPG*) (Hs00900358_m1); *TNFSF11* (*RANKL*) (Hs00243522_m1).

#### Mouse

*Cdkn2a* (Mm00494449_m1); *Gapdh* (Mm99999915_g1); *Hprt1* (Mm03024075_m1); *Il6* (Mm00446190_m1); *Tnfrsf11b* (*Opg*) (Mm01205928_m1); *Tnfsf11* (*Rankl*) (Mm00441906_m1).

The following primer pairs were additionally used:

- Cdkn1a Forward: 5′- CCT CCC AAG ATA GCC GAG TT -3′ Reverse: 5′- AGA CGA CAC AGG TGA GGA AG -3′
- *Il1β* Forward: 5′- ACA AGG AGA ACC AAG CAA CG -3′ Reverse: 5′- GGG TGT GCC GTC TTT CAT TA -3′
- *Tnf* Forward: 5′- CAG GCG GTG CCT ATG TCT C -3′ Reverse: 5′- CGA TCA CCC CGA AGT TCA GTA G -3′

SYBR Green-based reactions were performed Power SYBR® Green PCR Master Mix (Applied Biosystems) or Platinum® SYBR® Green qPCR SuperMix-UDG with ROX (Invitrogen, Carlsbad, CA, USA).

### 2.15 RNA Sequencing and comparison

Total RNA from MC3T3-E1 cells doxorubicin-treated or untreated was extracted on day 7 as described above. Quality assessment, library preparation, RNA sequencing and bioinformatic analysis was carried out by Novogene (Novogene GmbH, Munich, Germany). Libraries were sequenced using an Illumina NovaSeq X Plus Series (PE150) system. RNA sequencing data was compared to transcriptomic data of humeri from young (2 months) and aged (30 months) old male mice published by Kaya et al. (Kaya, Schurman, Dole, Evans, & Alliston, 2022) accessible on https://www.mouse2human.org.

### 2.16 Statistical Analysis

Data were analyzed using GraphPad Prism version 11.0.2 (100) for macOS (GraphPad Software, Boston, MA, USA, www.graphpad.com). Datasets with n ≤ 5 were tested for outliers by means of the Grubbs’ outlier test. Outliers were not included in the statistical analyses. Each datapoint represents an independent biological replicate (donor or experiment for MC3T3-E1 cells). Data sets of two groups were analyzed by means of two-tailed (multiple) t-test. Data sets of three groups were analyzed by means of ordinary one-way ANOVA with post-hoc Dunnett’s multiple comparison. In each case, significance level was set to α = 0.05.

## 3. Results

### 3.1 Doxorubicin induces a stable SIPS phenotype in MC3T3-E1 cells

To enable an in-depth characterization of senescent OBs, two commonly used models were compared to identify a stable and reproducible *in vitro* model of stress-induced premature senescence (SIPS). Doxorubicin has previously been described as a potent inducer of cellular senescence in human chondrocytes (Kirsch et al., 2022). Based on these findings, a concentration of 0.1 μM doxorubicin was used to test for induction of senescence in MC3T3-E1 cells. In addition, H_2_O_2_ treatment was applied as an alternative senescence-inducing stimulus based on prolonged oxidative stress exposure, as previously described (Tripathi, Yen, & Singh, 2020). To elucidate the induction of SIPS, senescence-associated β-galactosidase (SA-β-gal) staining revealed a pronounced increase to 87.1% positive cells following doxorubicin treatment. In contrast, after H_2_O_2_ treatment only a modest increase of 12.5% was observed (Fig. 1A, B). Gene expression analysis of senescence-associated cell cycle regulators demonstrated significantly increased expression of cyclin-dependent kinase inhibitor 1 (*Cdkn1a*) and cyclin-dependent kinase inhibitor 2a (*Cdkn2a*) in doxorubicin-treated cells, whereas H_2_O_2_ treatment had no significant effect compared to untreated cells (Fig. 1C). Ki-67 immunofluorescence staining demonstrated reduced cellular proliferation following both doxorubicin (-73.1%) and H_2_O_2_ (-26.0%) treatment compared to untreated cells (Fig. 1D, E). Notably, approximately 40% of H_2_O_2_-treated cells remained positive for Ki-67, compared with only 2.2% of doxorubicin-treated cells. In addition to the marked decline in Ki-67 expression, cell cycle analysis using Vybrant^TM^ staining was performed to determine the phase at which cell cycle arrest occurred following doxorubicin treatment. The cell cycle arrest in doxorubicin-treated cells was characterized by a shift from the G_0_/G_1_ phase towards G_2_/M phase. Accordingly, the G_0_/G_1_ population was reduced by 32.9%, while the G_2_/M phase was increased by 37.6% compared to untreated control (Fig. 1F). Expression levels of SASP-associated cytokines tumor necrosis factor (*Tnf*; 6.8-fold) and interleukin 6 (*Il6*; 4.6-fold) were significantly induced by doxorubicin, but not by H_2_O_2_. Analysis of osteoclastogenesis regulators revealed a significant increase for receptor activator of nuclear factor kappa-B (*Rankl*; 6.7-fold), but no change in case of osteoprotegerin (*Opg*; 1.8-fold) in doxorubicin-treated cells. (Fig. 1G). Calculation of the *Rankl*/*Opg* ratio (Fig. 1H) revealed a significant increase in doxorubicin-treated cells, whereas no significant shift was observed following H_2_O_2_ treatment, suggesting that senescent osteoblasts promote a pro-osteoclastogenic microenvironment. As cellular senescence is primarily driven by DNA damage γ-H2A.X staining was performed to confirm persistent DNA damage. Doxorubicin-treated cells exhibited an increase in γ-H2A.X-positive nuclei of 63.1% compared to untreated control, indicating persistent double-strand DNA break formation (Fig. 1I, J). In contrast, γ-H2A.X was increased 30 min after first time H_2_O_2_ administration but returned to baseline levels after 24 h of H_2_O_2_ exposure (Fig. 1J).

**Figure 1:**
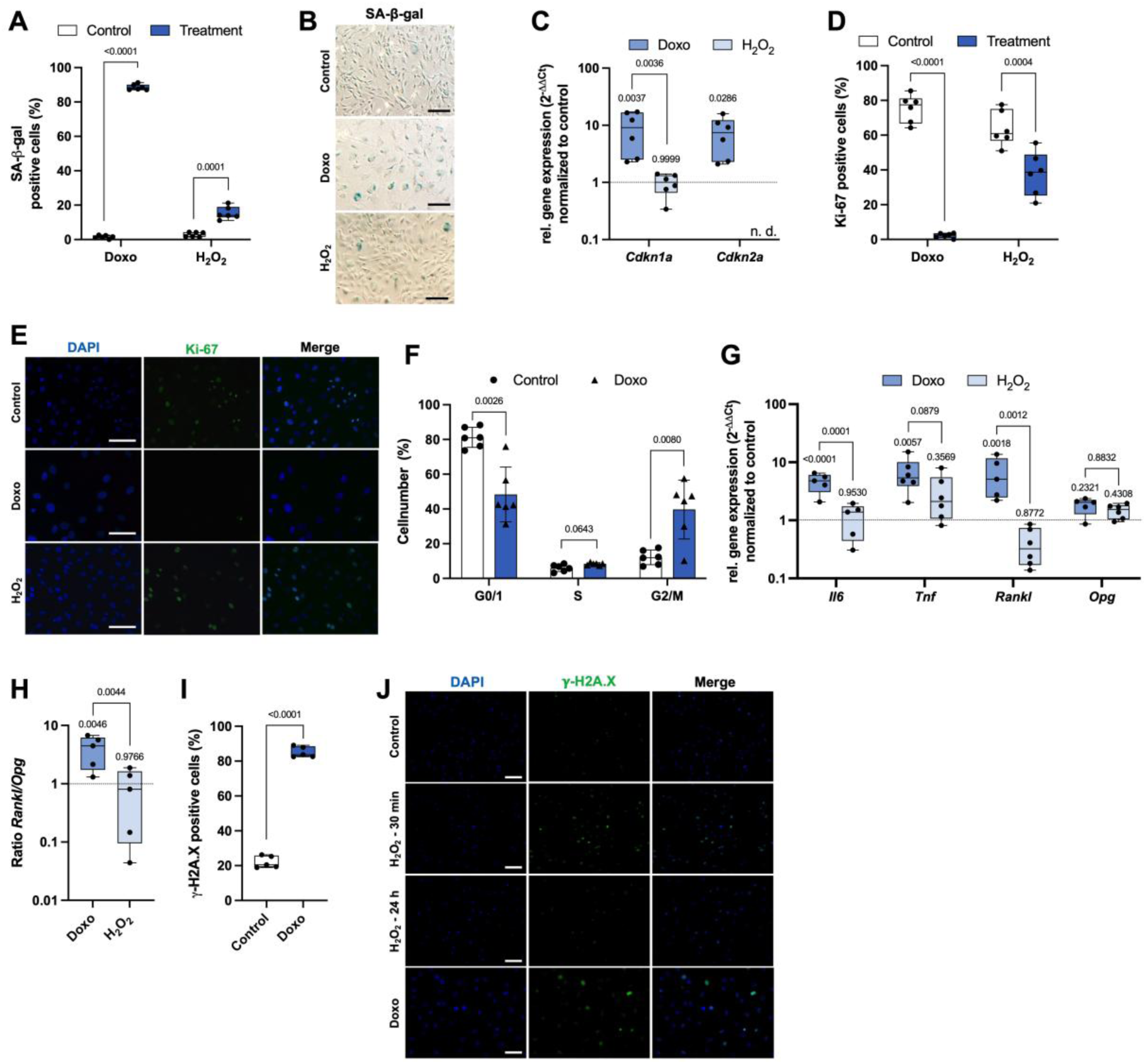
Establishment and validation of stress-induced premature senescence in MC3T3-E1 cells. (A) Quantification and (B) exemplary of SA-β-gal staining of doxorubicin- or H_2_O_2_-treated MC3T3-E1 cells, Scalebar = 150 µm. (C) Gene expression levels of *Cdkn1a* and *Cdkn2a* were analyzed of doxorubicin- or H_2_O_2_-treated cells. (D) Quantification and (E) representative immunofluorescence images of Ki-67 staining, Scalebar = 100 µm. (F) cell cycle analysis of doxorubicin-treated cells. (G) Gene expression levels of SASP markers *Il6*, *Tnf*, and osteoclastogenesis regulators *Rankl* and *Opg*. (H) *Rankl*/*Opg* ratio of MC3T3-E1 cells treated with doxorubicin or H_2_O_2_. (I) Quantification and (J) representative immunofluorescence staining for γ-H2A.X stained MC3T3-E1 cells treated with 100 µM H_2_O_2_ for 30 min and 24 h and day 7 of 0.1 µM doxorubicin-treated cells, Scalebar = 100 µm. Data are presented as box plot with median, whiskers min to max. Significant differences between groups are indicated. Statistical analysis: (A, D, F, I) multiple paired t-tests; (C, G, H) ordinary one-way ANOVA for each gene or treatment with post-hoc Dunnetts’ multiple comparisons test with control and doxorubicin-treated cells set as “controls” to compare the groups. Control = untreated cells; Doxo = Doxorubicin. n ≤ 5.

Only the doxorubicin-based SIPS *in vitro* model was used in subsequent experiments to induce senescence in osteoblasts, as this model showed robust expression of senescence-associated markers.

### 3.2 Senescence impairs Osteogenic Differentiation Capacity in MC3T3-E1 cells

To investigate the impact of senescence on osteogenic differentiation, untreated (non-senescent) and doxorubicin-treated (senescent) MC3T3-E1 cells were cultured under osteogenic conditions and analyzed at early (day 7), intermediate (day 14), and late (day 21) stages of differentiation. Qualitative and quantitative alkaline phosphatase (ALPL) staining revealed a significant reduction in ALPL activity in senescent cells at day 7 compared to non-senescent (Fig. 2A, B). Furthermore, Alizarin Red S staining demonstrated significantly reduced calcium deposition in senescent cells at day 14 (Fig. 2C, D) and day 21, indicating impaired matrix mineralization and reduced osteogenic differentiation capacity (Fig. 2E, F).

**Figure 2:**
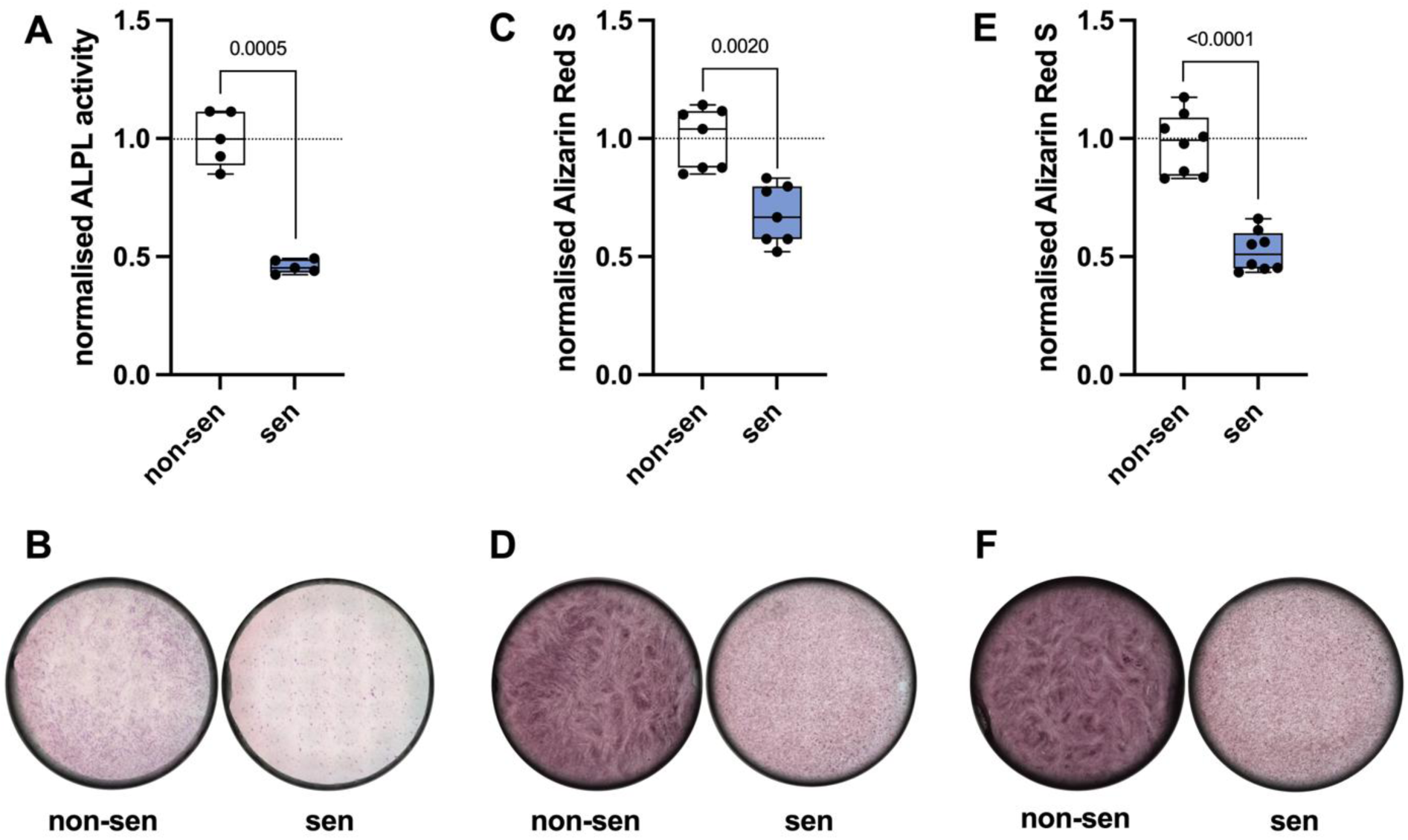
Osteogenic differentiation capacities of senescent MC3T3-E1 cells. (A) Quantified ALPL activity of early (day 7) osteogenic differentiated non-senescent and senescent MC3T3-E1 cells and (B) exemplary qualitative ALPL staining of non-senescent and senescent cells. (C) Quantified calcium deposition by Alizarin Red S staining on day 14 of differentiation and (D) exemplary staining images of non-senescent and senescent cells. (E) Quantification of Alizarin Red S staining on day 21 of osteogenic differentiation and (F) representative images of non-senescent and senescent MC3T3-E1 cells. Data are presented with box plots with median, whiskers min to max. Significant differences between groups are indicated. Statistical analysis: (A, C, E) paired t-test. Non-sen = non-senescent cells (untreated control), sen = senescent cells (doxorubicin-treated). n ≤ 5.

### 3.3 Cellular Senescence Alters Mitochondrial Physiology in MC3T3-E1 cells

Given the central role of the SAMP in the promotion and maintenance of cellular senescence, mitochondrial function and morphology were investigated in greater detail. Senescent MC3T3-E1 cells exhibited significantly reduced mitochondrial membrane potential, as indicated by a decreased red-to-green fluorescence ratio in the JC-1 assay (Fig. 3A) and increased mitochondrial superoxide accumulation compared to non-senescent controls (Fig. 3B). This was accompanied by pronounced mitochondrial elongation, as revealed by transmission electron microscopy (TEM) analysis (Fig. 3C). Mitochondrial bioenergetics were further investigated using Seahorse extracellular flux analysis. Senescent cells exhibited reduced basal oxygen consumption rate (OCR; Fig. 3D), while the overall proton efflux rate (PER; Fig. 3E) was also decreased. In addition, ATP-linked OCR (Fig. 3F), ATP production by glycolysis (ATPgly; Fig. 3G) and ATP production by oxidative phosphorylation (ATPox; Fig. 3H), were significantly reduced, indicating a senescence-associated bioenergetic impairment characterized by diminished ATP-production capacity and reduced metabolic flexibility (Fig. 3I).

**Figure 3:**
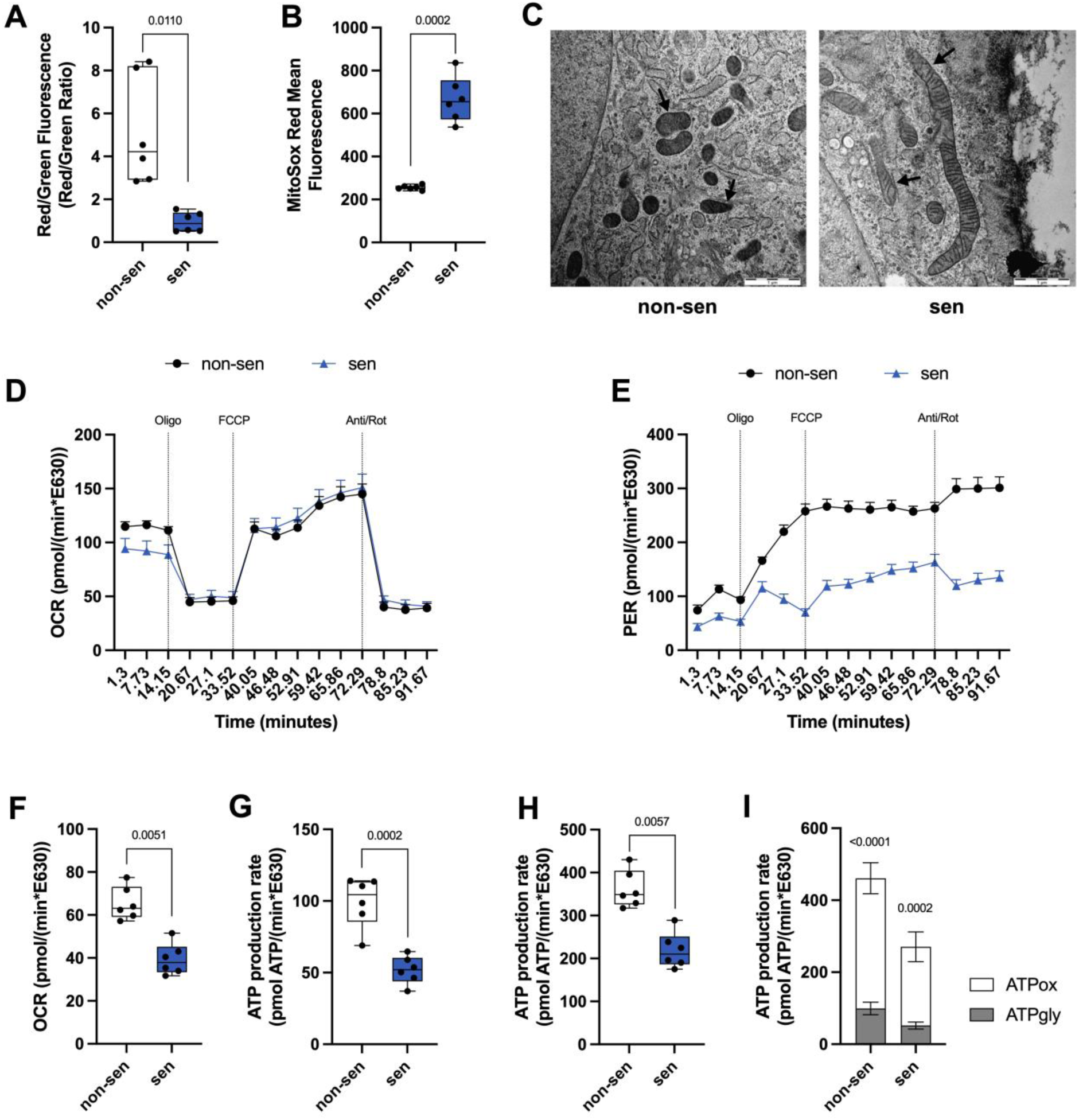
Characterization of senescence-associated mitochondrial phenotype in MC3T3-E1 cells. (A) Mitochondrial membrane potential. (B) Super oxide accumulation in non-senescent and senescent cells. (C) Exemplary TEM images of mitochondria of non-senescent and senescent MC3T3-E1 cells, mitochondria indicated with black arrows, scalebar = 1 µm. (D) OCR, (E) PER, (F) ATP-linked OCR, (G) ATP produced by glycolysis (ATPgly) and (H) ATP produced by oxidative phosphorylation (ATPox) of non-senescent and senescent cells. (I) Summary of ATPgly and ATPox. Data are presented with box plots with median, whiskers min to max and time-course line plot. Significant differences between groups are indicated Statistical analysis: (A, B, F, G, H) paired t-test, (I) multiple paired t-test. Non-sen = non-senescent cells (untreated control), sen = senescent cells (doxorubicin-treated). n = 6.

### 3.4 Doxorubicin-induced SIPS in MC3T3-E1 cells mirrors key molecular features of skeletal aging in vivo

To assess whether the *in vitro* doxorubicin-based SIPS model in MC3T3-E1 recapitulates transcriptional features of aged bone *in vivo*, we compared our dataset with a published transcriptomic dataset of young and aged mouse humeri as previously published (Kaya et al., 2022). A principal component analysis (PCA) of the *in vitro*-generated data revealed a clear separation between senescent (doxorubicin-treated) and non-senescent (vehicle-treated) MC3T3-E1 cells, indicating distinct global transcriptional profile associated with cellular senescence (Fig. 4A). By differential gene expression analysis, we could identify a total of 708 downregulated genes and 1331 upregulated genes when comparing senescent and non-senescent MC3T3-E1 (Fig. 4B). Functional enrichment analysis using Metascape revealed significant enrichment of genes associated with cell cycle regulation and chromosome dynamics, as well as DNA-associated process (e.g. DNA replication) (Fig. 4C). These findings are consistent with the established mechanism of doxorubicin as a DNA-damaging agent, also reflected by the experimental observations of increased γ-H2A.X positive cells, altered cell-cycle distribution, and elevated expression of the senescence-associated cell-cycle regulators *Cdkn1a* and *Cdkn2a*. Together, these findings suggest that persistent DNA damage response pathways contribute to the establishment and maintenance of the senescent phenotype in the doxorubicin-based *in vitro* model.

**Figure 4:**
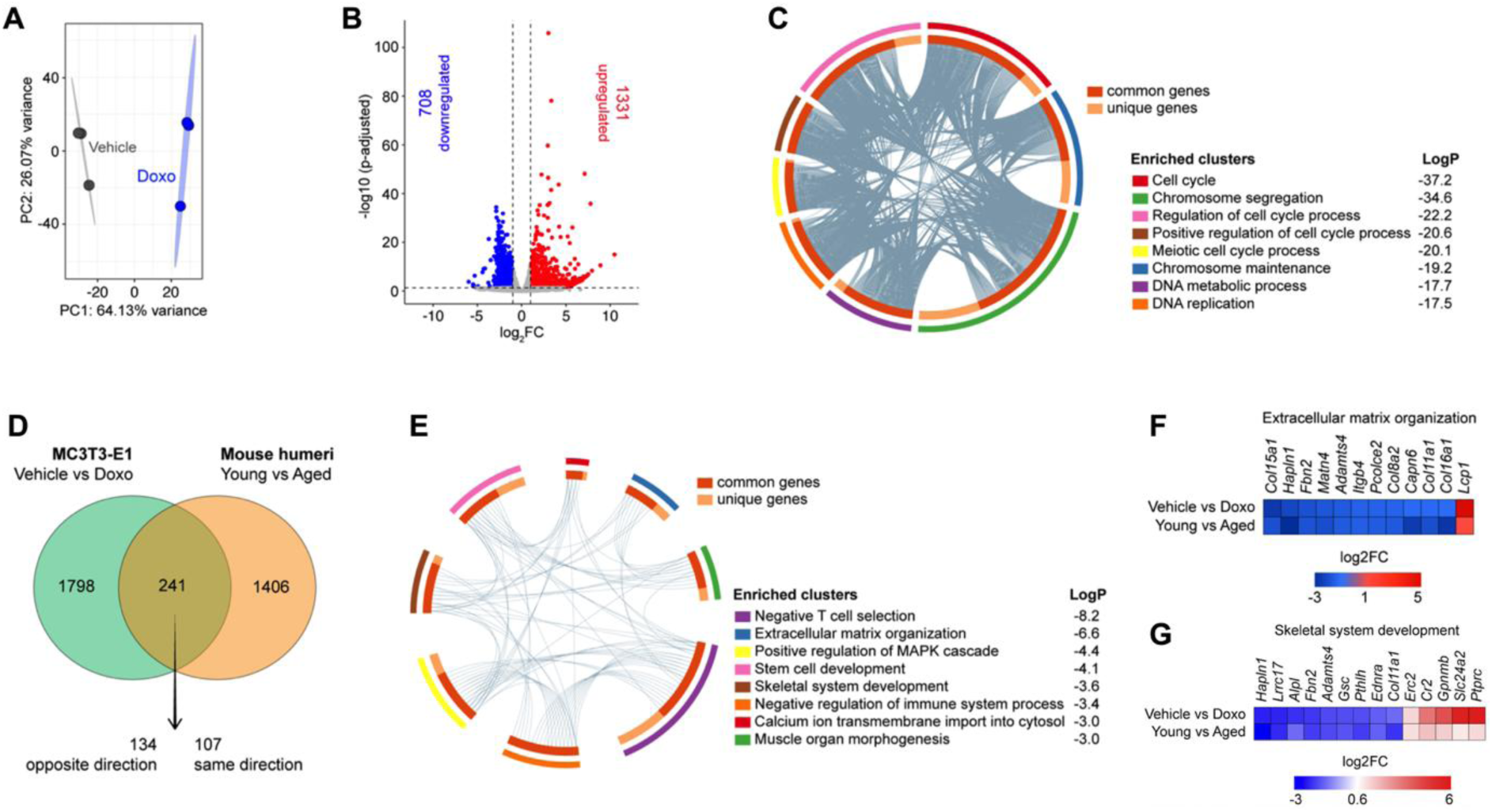
Comparative transcriptomic analysis of in vitro and in vivo bone senescence models. (A) Principal component analysis (PCA), (B) volcano plot and (C) metascape of senescent (doxorubicin-treated) and non-senescent (vehicle-treated) MC3T3-E1 cells. (D) Comparison between the transcriptomic data set of the *in vitro* SIPS model and aged murine humeri. Corresponding (E) metascape and (F, G) heatmap of selected clusters. n=3.

Comparing published transcriptomic dataset of young and aged mouse humeri (Kaya et al., 2022) with our data identified 241 overlapping differentially expressed genes (Fig. 4D). While 107 genes were regulated in the same direction, 134 exhibited opposite regulation, indicating partial transcriptional overlap with skeletal aging *in vivo*. Metascape revealed significant enrichment associated with e.g., extracellular matrix organization, positive regulation of MAPK cascade, stem cell as well as skeletal system development, negative regulation of immune system process, and calcium ion transmembrane import into cytosol (Fig. 4E).

Because extracellular matrix organization represents a key determinant of bone quality and osteoblast function, this pathway was investigated in greater detail. Several genes involved in matrix composition, organization, and collagen homeostasis, including collagen type XV alpha 1 chain (*Col15a1*), fibrillin 2 (*Fbn2*), procollagen C-endopeptidase enhancer 2 (*Pcolce2*), and multiple collagen family members, were similarly downregulated in both datasets. In contrast, lymphocyte cytosolic protein 1 (*Lcp1*) was identified as commonly upregulated gene across both datasets (Fig. 4F). Besides extracellular matrix organization, enrichment analysis also highlighted pathways related to skeletal system development (Fig. 4G). Multiple genes related to skeletal system development, osteogenic differentiation and bone homeostasis including, leucine rich repeat containing 17 (*Lrrc17*), alkaline phosphatase (*Alpl*) and hyaluronan and proteoglycan link protein 1 (*Hapln1*) were similarly down regulated in both datasets. In contrast genes involved in immune regulatory processes, tissue remodeling, and calcium homeostasis including, complement receptor 2 (*Cr2*), transmembrane glycoprotein nmb (*Gpnmb*), and solute carrier family 24 member 2 (*Slc24a2*) were commonly upregulated in both datasets. Overall, the dysregulation of genes related to extracellular matrix organization and skeletal development aligns with impaired *in vitro* osteogenic differentiation of senescent MC3T3-E1 cells and the characteristic decline of osteoanabolism in the aged skeleton.

### 3.5 Impaired osteogenic function and mitochondrial dysfunction are conserved in human primary osteoblasts

The transcriptomic overlap with aged murine bone suggested that the established model recapitulates selected features of skeletal aging. To test for translational relevance and to determine whether key senescence-associated cellular and mitochondrial alterations are conserved across species, also primary human osteoblasts (hOB) were treated with doxorubicin as described above and the level of senescence determined. To assess persistent DNA damage and exclude pre-existing DNA damage associated with donor age, γ-H2A.X immunofluorescence staining was performed. Following senescence induction, hOB exhibited an increase of 73.8% in γ-H2A.X-positive nuclei compared to untreated controls (8.14% γ-H2A.X-positive nuclei), indicating elevated double-strand DNA breaks (Fig. 5A, B). Cell cycle analysis using Vybrant^TM^ staining revealed an accumulation of cells in the G_0_/G_1_ phase, with an increase of 4.5% following senescence induction. In contrast, the S phase population decreased by 1.8%, while the G_2_/M population showed a significant decrease of 6.6% (Fig. 5C). Ki-67 immunofluorescence staining revealed a significant decrease in Ki-67-positive cells by 28.6% following senescence induction (Fig. 5D, E). Gene expression analysis of senescence associated cell cycle regulators demonstrated a 9.2-fold increase in *CDKN1A* levels, whereas only a 2.8-fold increase for *CDKN2A* could be observed (Fig. 5F). Similarly, the proportion of SA-β-gal positive cells was elevated by 68.3% compared to untreated controls (Fig. 5G, H). Analysis of SASP-associated cytokines revealed an induction in interleukin 1β (*IL1β*; 15.1-fold), interleukin 6 (*IL6*; 3.5-fold), interleukin 8 (*IL8*; 4.8-fold), and tumor necrosis factor (*TNF*; 10.4-fold) expression following doxorubicin treatment compared to untreated controls. To investigate the influence of senescence on osteoclastogenesis-associated regulators, expression of receptor activator of nuclear factor kappa-B ligand (*RANKL*) and osteoprotegerin (*OPG*) was assessed. Senescence induction resulted in significantly increased *RANKL* (18.5-fold) expression, whereas no change was observed in case of *OPG* (0.95-fold) (Fig 5I). Calculation of the *RANKL*/*OPG* ratio demonstrated a significant increase about 12-fold in senescent hOBs, indicating a senescence-associated shift towards an osteoclastogenic microenvironment (Fig. 5J). Osteogenic differentiation for 21 days revealed a significant decrease in calcium deposition in senescent hOB compared to the control (Fig. 5K, L).

**Figure 5:**
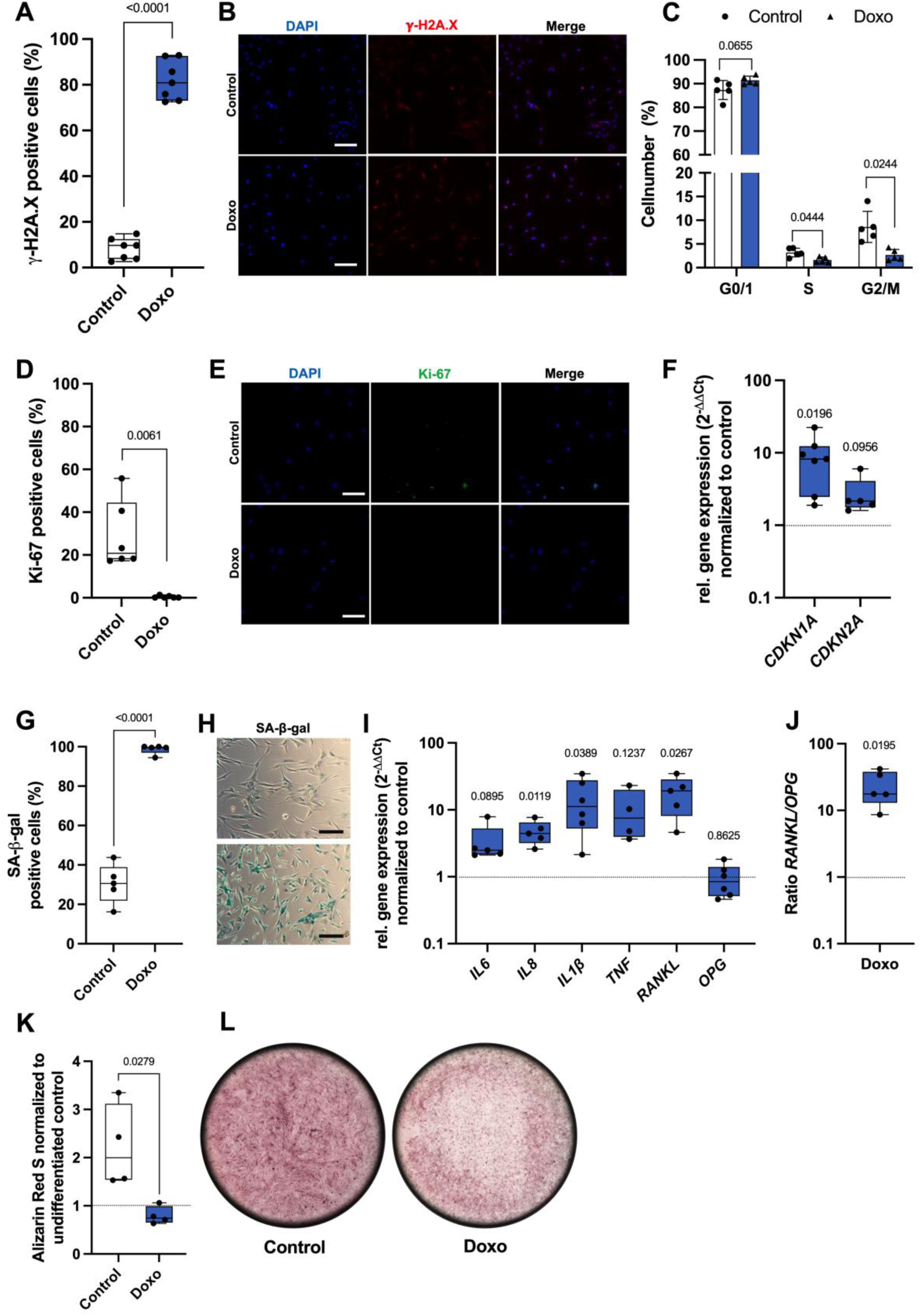
Validation of stress-induced premature senescence in primary human osteoblasts. (A) Quantification and (B) exemplary imaged of immunofluorescence staining of γ-H2A.X and (C) cell cycle analysis of doxorubicin treated hOB, scalebar = 100 µm. (D) Quantification and (E) immunofluorescence Ki-67 staining, scalebar = 100 µm. (F) Gen expression levels of *CDKN1A* and *CDKN2A*. (G) Quantification and (H) brightfield images of SA-β-gal staining, scalebar = 150 µm. (I) Gene expression analysis of SASP markers and osteoclastogenesis regulators. (J) *RANKL*/*OPG* Ratio of doxorubicin treated hOB. (K) Alizarin Red S staining quantified and (L) exemplary whole-well images of control and Doxo on day 21 of osteogenic differentiation. Data are presented with box plots with median, whiskers min to max. Significant differences between groups are indicated. Statistical analysis: (A, D, G, J, K) paired t-test, (C, F, I) multiple paired t-test. Control = untreated, Doxo = doxorubicin-treated. n ≤ 4.

As doxorubicin treatment of hOB resulted in the establishment of a stable senescent phenotype, including a SASP, untreated cells are hereafter designated as non-senescent, whereas doxorubicin-treated cells are referred to as senescent.

Mitochondrial function was further analyzed using Seahorse Flux Analyzer. Senescent hOB showed a reduced basal OCR (Fig. 6A) and overall PER (Fig. 6B). Moreover, senescence was associated with a reduction in ATP-linked OCR (Fig. 6C), glycolytic ATP production (ATPgly; Fig. 6D) as well as oxidative ATP production (ATPox; Fig. 6E), and a diminished maximal respiration. Overall, Seahorse analysis indicated impaired respiratory reserve capacity and a senescence-associated bioenergetic defect, characterized by reduced ATP production from both glycolytic and oxidative pathways. Additionally, TEM analysis revealed a pronounced elongation of mitochondrial morphology and accumulation of autophagosomes in senescent hOB (Fig. 6F).

**Figure 6:**
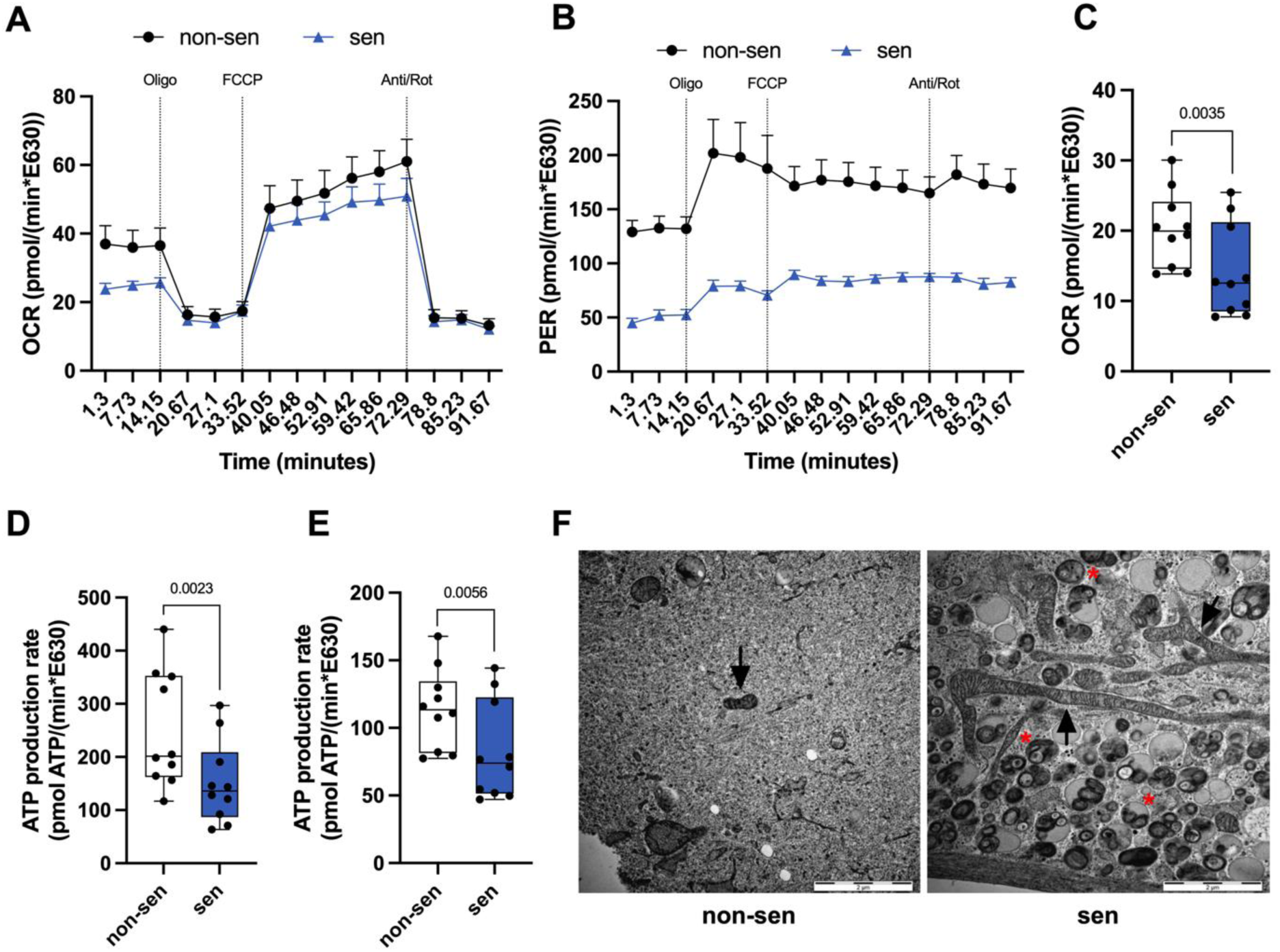
Influence of cellular senescence on bioenergetic processes of hOB. (A) Oxygen consumption rate (OCR), (B) proton efflux rate (PER), (C) ATP-linked OCR, (D) ATP produced by glycolysis (ATPgly) and (E) ATP produced by oxidative phosphorylation (ATPox) of control and doxorubicin treated hOB. (F) Exemplary TEM images of non-senescent and senescent hOB, mitochondria are indicated by black arrows, autophagosomes by red asteriks, Scalebar = 2 µm. Data are presented with box plots with median, whiskers min to max, grouped interleaved scatter with bars and time-course line plot. Significant differences between groups are indicated. Statistical analysis: (C, D, E) paired t-test. Non-sen = non-senescent (untreated hOB), sen = senescent (doxorubicin treated hOB). n ≤ 3.

## 4. Discussion

Over the past few years cellular senescence has been recognized as a central hallmark of skeletal aging and age-related bone disease, such as osteoporosis (Riegger et al., 2023). Senescent osteoblast lineage cells lose their regenerative and matrix-forming capacity (Kasama et al., 2023; Pignolo et al., 2021). Since osteoblast differentiation and mineralization are essential for maintaining skeletal homeostasis, accumulation of senescent osteoblast may directly contribute to the age-associate decline in bone formation observed during osteoporosis and skeletal ageing (Khosla, Farr, & Monroe, 2022). Moreover, the release of pro-inflammatory SASP factors could further compromise osteoanabolism in non-senescent bone cells in a paracrine manner, as previously reported in human chondrocytes (Maurer, Kirsch, Ruths, Brenner, & Riegger, 2025).

In this study, we established and characterized a robust *in vitro* model of SIPS in murine MC3T3-E1 cells by comparing two different inducers with distinct modes of action: DNA damage by doxorubicin and prolonged oxidative stress by hydrogen peroxide. Our findings demonstrate that doxorubicin induces a stable senescent phenotype, characterized by irreversible proliferation arrest, increased SA-β-gal activity, altered cell cycle distribution, SASP factor expression, and impaired osteogenic differentiation, in both MC3T3-E1 cells and hOBs. Osteoblast function was highly impaired upon senescent induction. Doxorubicin-treated MC3T3-E1 cells exhibited significantly reduced alkaline phosphatase activity and calcium deposition during osteogenic differentiation.

In contrast, H_2_O_2_ treatment induced senescence in only a subset of MC3T3-E1 cells, indicating a heterogenous response that manifested as relatively subtle changes under the selected experimental conditions. Oxidative stress-induced senescence is strongly dependent on dose, exposure time, and cellular recovery conditions. Therefore, transient ROS exposure may elicit a partial or less stable senescence-like phenotype (Jin et al., 2024; Robert, Kennedy, & Crasta, 2024). Moreover, the high proliferative nature of MC3T3-E1 cells may have obscured SASP-associated changes, suggesting that H_2_O_2_-based senescence induction may be less suitable for rapidly dividing cell models. In depth analyses on mitochondrial properties in doxorubicin treated MC3T3-E1 as well as primary human OBs revealed a remarkable SAMP, including functional and morphological alternations. Moreover, comparison of senescent MC3T3-E1 cells with aged murine bone cells confirmed a partial overlap with *in vivo* aging signatures, altogether supporting its translational relevance of the established *in vitro* senescence model. Doxorubicin treated MC3T3-E1 cells might thus serves as model system for senescence in primary human OBs or murine bone cells.

Mechanistically, doxorubicin treatment resulted in the expected persistent γ-H2A.X formation, confirming accumulation of DNA double-strand breaks in the treated cells (Swift, Rephaeli, Nudelman, Phillips, & Cutts, 2006). Persistent DNA damage is considered a major driver of senescence maintenance through activation of the DNA damage response (DDR), ultimately leading to stable cell cycle arrest mediated by p21 (*CDKN1A*) and p16 (*CDKN2A*) signaling pathways (Campisi & d’Adda di Fagagna, 2007). Similarly, we could observe increased *Cdkn1a* and *Cdkn2a* expression for MC3T3-E1 cells alongside altered cell cycle distribution characterized by accumulation in G_2_/M phase. In contrast, doxorubicin induced senescence in hOB exhibited an accumulation in G_0_/G_1_ phase, as we previously observed in human chondrocytes (Kirsch et al., 2022). Although, senescence has commonly been associated to the G_1_-phase of the cell cycle, it can also occur in G_2_ (Kumari & Jat, 2021; Valentijn, Falke, Nguyen, & Goldschmeding, 2018). Although H_2_O_2_-induced stress-induced premature senescence in a well-established model has been successfully applied in numerous cell types (Duan, Duan, Zhang, & Tong, 2005; Kiyoshima et al., 2012), its efficiency is highly dependent on factors including cell type, oxidant concentration, exposure duration, and recovery conditions. The relatively modest and heterogenous response observed in MC3T3-E1 cells under our experimental conditions therefore likely reflects the specific characteristics of this cell line rather than a general limitation of oxidative stress-induced senescence. These findings highlight the importance of validating senescence induction protocols for each experimental model and support the use of doxorubicin as the more robust approach for inducing stable senescence in MC3T3-E1 osteoblasts.

In addition to impaired osteogenic differentiation, senescent osteoblasts exhibited an increased *RANKL*/*OPG* ratio, indicating a shift towards a more pro-osteoclastogenic phenotype. Given the central role of the *RANKL*/*OPG* axis in regulating osteoclast differentiation and activation (Udagawa et al., 2021), these findings suggest that senescent osteoblasts may contribute to age-related bone less not only through diminished bone formation but also by actively promoting bone resorption. Such a dual effect is particularly relevant in the context of osteoporosis, where impaired osteoblast function and enhanced osteoclast activity together drive progressive deterioration of bone mass and quality. This interpretation is supported by *in vivo* evidence demonstrating that selective elimination of senescent osteoblasts in aged mice reduced osteoclast numbers and attenuated age-related bone loss (Farr et al., 2017). Furthermore, conditioned medium derived from senescent cells has been shown to promote osteoclastogenesis *in vitro*, highlighting the importance of senescence-associated paracrine signaling in regulating bone remodeling (Farr et al., 2017)

A major finding of the present study is the pronounced SAMP observed in senescent osteoblasts. Senescent MC3T3-E1 exhibited reduced mitochondrial membrane potential and increased mitochondrial superoxide accumulation. In addition to reduced basal OCR, both senescent MC3T3-E1 cells and primary human osteoblasts exhibited lower PER together with reduced glycolytic (ATPgly) and oxidative phosphorylation-derived (ATPox) ATP production. Taken together, these findings suggest a global downshift in cellular energy metabolism rather than a specific defect in ATP synthesis, most likely reflecting combined impairments in electron transport, proton gradient maintenance and overall metabolic flux. Furthermore, mitochondrial dysfunction is increasingly recognized as both a driver and consequence of cellular senescence. Elevated mitochondrial ROS production can reinforce DNA damage and SASP signaling, thereby contributing to maintenance of the senescent phenotype (Miwa, Kashyap, Chini, & von Zglinicki, 2022), while persistent DNA damage itself may impair mitochondrial homeostasis through altered mitophagy, mitochondrial biogenesis and metabolic rewiring (Hruby & Higuchi-Sanabria, 2025). In line with this concept, the observed increase in mitochondrial superoxide accumulation together with reduced membrane potential supports the existence of a self-amplifying senescence-mitochondrial dysfunction axis in osteoblasts (Xiong, Guo, & Luo, 2025).

To further assess the translational relevance of our model, we compared the transcriptomic profile of senescent MC3T3-E1 cells with a published data set of young and aged murine humeri (Kaya et al., 2022). Despite the inherent differences between an *in vitro* model of stress-induced premature senescence and physiological skeletal aging *in vivo*, both datasets shared alterations in pathways associated with extracellular matrix organization, skeletal development, and cellular signaling. In particular, genes involved in extracellular matrix organization exhibited expression patterns similar to those previously reported in the transcriptomic analysis of flushed tibiae from aged mice (Nandy et al., 2024). Nandy and colleagues demonstrated that skeletal aging is accompanied by dysregulation of genes controlling collagen synthesis, extracellular matrix organization and matrix remodeling, highlighting progressive impairment of matrix homeostasis in bone. Notably, although Kaya et al. and Nandy et al. analyzed different stages of murine skeletal aging (30 and 22 months, respectively), the consistent enrichment of extracellular matrix-related pathways across both datasets suggests that these alterations are conserved throughout skeletal aging. The overlap observed in the present study therefore suggests that senescent osteoblasts recapitulate key molecular features of the aging bone microenvironment. Given the central role of osteoblasts in extracellular matrix synthesis and maintenance, these findings further support the concept that osteoblast senescence contributes to age-associated deterioration of bone quality and function.

Overall, the key hallmarks of cellular senescence identified in MC3T3-E1 cells were reproduced in primary human osteoblasts following doxorubicin treatment, demonstrating that the established model captures conserved features of osteoblast senescence across species. Although species-specific differences were observed, including the expression of *Cdkn2a*/*CDKN2A* and the predominant phase of cell cycle arrest, these variations likely reflect intrinsic biological differences between murine and human osteoblasts rather than fundamentally district senescence programs. The conservation of the major senescence-associated characteristics supports the translational relevance of the model and reinforce the concept that osteoblast senescence contributes to age-associated disruption of bone remodeling.

Nevertheless, several limitations should be considered. Pharmacologically induced SIPS, particularly in immortalized MC3T3-E1 cells, cannot fully recapitulate the complexity of chronological skeletal aging or the interactions of osteoblasts with the bone microenvironment *in vivo*. Despite these limitations, the established model demonstrated a remarkable degree of biological conservation. Although MC3T3-E1 cells and primary human osteoblasts differ in their embryonic origin and physiological context with MC3T3-E1 cells being derived from calvarial bone formed through intramembranous ossification and human osteoblasts isolated from long bone undergoing endochondral ossification, both models exhibited remarkably similar senescence-associated characteristics following doxorubicin treatment. This finding is particularly noteworthy, as osteoblasts from different skeletal sides are known to display distinct phenotypic, transcriptomic, and functional properties, including differential responses to biochemical stimuli. Furthermore, the transcriptomic overlap between senescent MC3T3-E1 cells and aged murine bone further supports the biological and translational relevance of the established osteoblast senescence model. However, these transcriptomic analyses remain primarily descriptive, and future studies integrating proteomic and functional validation will be required to determine how these transcriptional alterations contribute to age-related skeletal dysfunction.

In conclusion, we demonstrate that senescent osteoblasts develop an inflammatory and pro-osteoclastogenic SASP together with a pronounced SAMP, identifying mitochondrial dysfunction as a key component of the metabolic reprogramming accompanying cellular senescence. Given the central role of mitochondria in osteoblast differentiation, matrix mineralization, and calcium homeostasis these alterations may directly contribute to the impaired osteogenic capacity observed in senescent cells (Shapiro, Risbud, & Landis, 2024; Suh & Lee, 2024). Furthermore, senescent osteoblasts exhibited impaired osteogenic function accompanied by transcriptional alterations in pathways related to extracellular matrix organization and skeletal development, partially recapitulating molecular signatures of skeletal aging in vivo. Collectively, these findings support an active role for osteoblast senescence in age-associated bone remodeling and establish doxorubicin-induced senescence as a robust and translationally relevant model for investigating the molecular mechanisms underlying skeletal aging and the development of future senescence-targeted therapeutic strategies.

Collectively, out findings support the concept that osteoblast senescence and mitochondrial dysfunction act in concert to drive skeletal aging and disrupt bone remodeling. These results further establish the close interplay between cellular senescence and mitochondrial dysfunction in osteoblast biology and highlight this senescence-mitochondrial axis as a potential therapeutic target for age-related bone disease.

## Acknowledgements

We would like to thank the ULMTeC Core Facility Extracellular Flux Analyzer of the Medical Faculty at Ulm University for providing support, expertise and instrumentation.

We would also like to thank Dr. Clarissa Read, Jana Apolloni and Reinhard Weih of the Central Facility for Electron Microscopy at Ulm University for their support, expertise and instrumentation.

## Permission statement

No material from other sources requiring permission for reproduction is included in this manuscript.

## Author contributions

TF: experiment conduction and data analysis, visualization, manuscript writing, JRK: funding acquisition, conceptualization, supervision, manuscript writing. HV, MA, YN, MV: experiment conduction and data analysis, review and editing of manuscript. AI: funding acquisition, review and editing of manuscript. SN, HG, AS: review and editing of manuscript.

## Data availability statement

Data are contained within the article.

