## Supplement for "Mitochondrial dysfunction and impaired osteogenic capacity define stress-induced osteoblast senescence"

### Supporting Information

Supplementary Figure 1

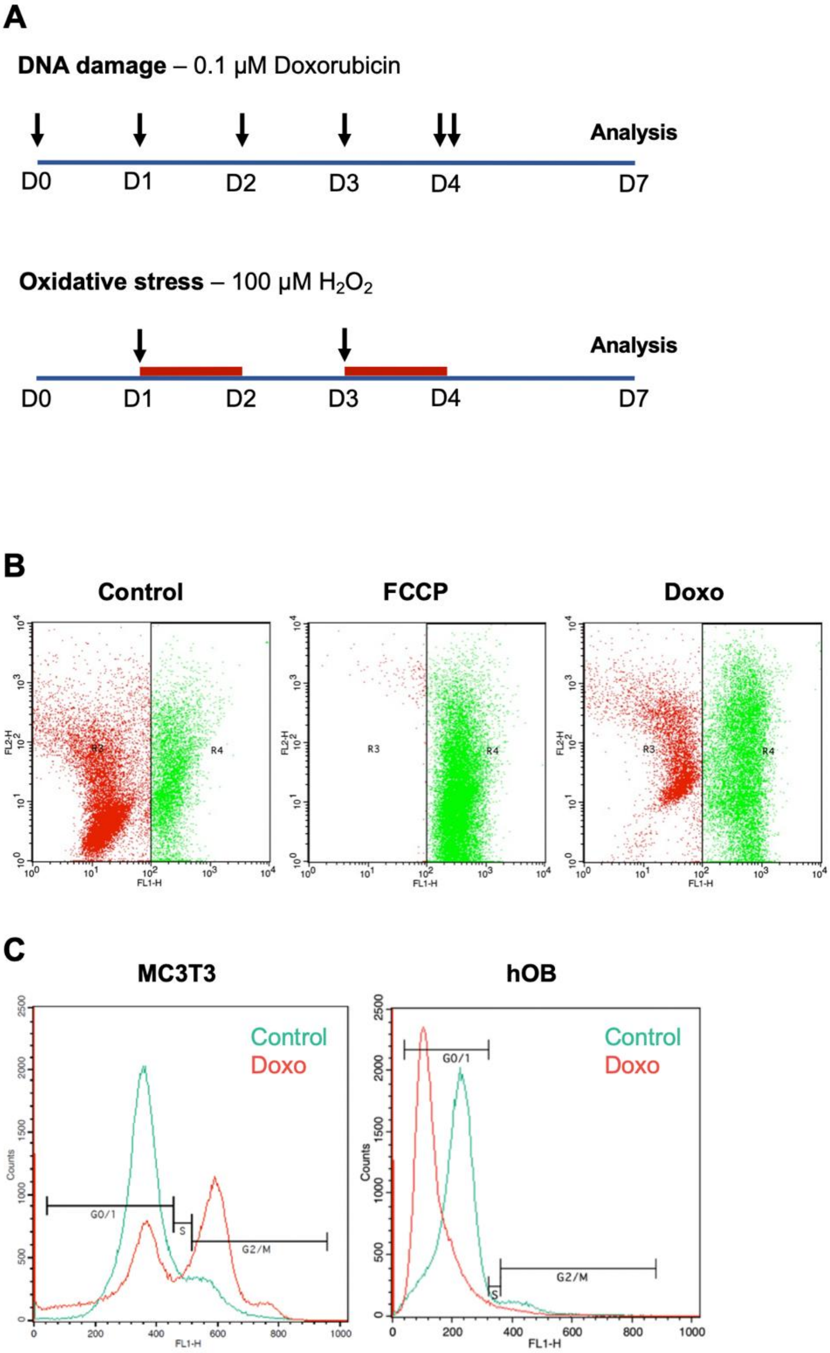


Supplementary Figure Legend

**Supplementary Figure 1: Stimulation scheme and flowcytometry gating strategies.** (A) Stimulation scheme for doxorubicin (Doxo) and $H_2O_2$ treatment. Black arrows indicating the addition of 0.1µM doxorubicin or 100 µM $H_2O_2$ respectively. Red bars indicating the 24h treatment duration for $H_2O_2$. (B) Flowcytometry gating strategy for JC-1 assay of MC3T3-E1 cells untreated and treated with doxorubicin, FCCP served as positive control. (C) Flowcytometry gating strategy for Cell cycle analysis by Vybrant$^{TM}$ cell cycle dye of untreated and doxorubicin-treated MC3T3-E1 and hOB.
